# Macrosystem community change in lake phytoplankton and its implications for diversity and function

**DOI:** 10.1101/2022.03.02.482597

**Authors:** Benjamin Weigel, Niina Kotamäki, Olli Malve, Kristiina Vuorio, Otso Ovaskainen

## Abstract

The combined effects of eutrophication, land-use and climate change are major threats to aquatic ecosystems, their biodiversity and integrity in sustaining ecosystem functions. Disentangling the mechanisms by which environmental change contributes to community assembly processes and species niches remains challenging, especially at macro-ecological scales. Here, we collated phytoplankton community data including 853 lakes along a 1200km latitudinal gradient, monitored over four decades, to quantify the spatio-temporal and scale-dependent environmental impacts on species niches and assembly processes while accounting for species traits and phylogenetic constraints. Our results demonstrate the emergence of novel and widespread community composition clusters in previously more uniform communities. While total species richness remained relatively stable, changes in community weighted mean traits of the clusters indicate functional differences. A robust phylogenetic signal of species responses to the environment indicates strong niche conservatism and low taxonomic dispersion. Our findings imply profound spatio-temporal structuring of species co-occurrence patterns and highlight emerging functional differences of lake phytoplankton communities to environmental change over space and time.

## INTRODUCTION

Freshwater ecosystems are biodiversity hotspots, vital for human well-being and provide a plethora of crucial ecological and socio-economic services (Falkenmark 2003; Myers 2003; Dudgeon *et al*. 2006). At the same time, lakes are among the most vulnerable ecosystems to anthropogenic pressures (Mammides 2020). At the terrestrial-aquatic interface, lakes are influenced by both, aquatic drivers such as water temperature, water inflow and mixing as well as by terrestrial impacts such as nutrient loadings from agriculture, soil composition from adjacent forests and fields, and urban areas in the lake catchment (Dudgeon *et al*. 2006). As primary producers, phytoplankton communities are directly and, due to their short generation time, rapidly affected by such impacts and mirror the ecological state of aquatic systems (Lepistö *et al*. 2004; Padisák *et al*. 2006). Recognizing lakes as sentinels for environmental impacts (Williamson *et al*. 2008), it is paramount to have a profound understanding of how species communities in lakes, such as phytoplankton, respond to anthropogenic changes at multiple spatial and temporal scales. Such knowledge supports the implementation of water management strategies such as the Water Framework Directive that aims at good ecological status of all surface waters in the European Union member states (European Comission 2000).

While lakes are often studied as closed and isolated environments at local or regional scales, they are often part of a connected network linked by watersheds and river basins (Soranno *et al*. 1999; Heino *et al*. 2020), enabling species dispersal and hierarchically structured impacts of environmental changes. It is well known that community dynamics and species diversity are scale dependent (Chase & Leibold 2002; Chase *et al*. 2019) and contingent on mechanisms such as competitive interactions, niche differentiation, and environmental filtering (Hardin 1960; Keddy 1992; Mayfield & Levine 2010). Despite recent research having highlighted lakes as meta-ecosystems at the waterscape level to account for these processes at different spatial scales (Heino et al 2020), knowledge about community assembly processes and the contributing abiotic and biotic interactions remains scarce. To shed further light on this, approaches for understanding community dynamics that include information on functional and evolutionary characteristics of species can be useful when investigating impacts on ecosystems, their biodiversity, and ecosystem function. Trait-based approaches have already proven beneficial in the assembly rule framework (McGill *et al*. 2006; Cadotte *et al*. 2015) since traits can explain species responses to environmental gradients (Edwards *et al*. 2013; Gagic *et al*. 2015). They have recently also been identified as promising diagnostic tool in water management planning, where they can aid to explain the underlying mechanisms and reasons for not achieving good ecological status under multiple environmental pressures (Baattrup-Pedersen *et al*. 2019; Carvalho *et al*. 2019). Hence, some traits may be good predictors for species occurrences considering e.g. their environmental niche, resource competition or dispersal, while other traits may be more relevant in the light of ecosystem services and water quality management, such as toxicity of occurring species (Grizzetti *et al*. 2019). However, certain traits may not only be linked to species that have undergone environmental filtering, but also to evolutionary constraints of closely related species being likely to share similar traits and occurrence patterns. This phenomenon is known as phylogenetic niche conservatism (Harvey & Purvis 1991).

Recent advances in statistical modelling frameworks in community ecology enable us to quantify how species responses to environmental covariates depend on species traits and phylogenetic relationships (Ovaskainen & Abrego 2020). Such tools pave new avenues to disentangle the scale dependent community assembly processes and their implications for diversity and function. Here we use an extensive phytoplankton community data set from the Finnish national lake monitoring program, including 853 lakes spanning the whole country of Finland, and sampled over four decades (1977-2017). We model species community composition at the sample level, highlight environmental filtering by quantifying species-specific variance partitioning of environmental covariates, and determine species co-occurrence patterns at different hierarchical spatial and temporal levels. To highlight emerging differences in community composition over space and time, we use a clustering algorithm to uncover different regions of common community profiles. Incorporating six traits, reflective of resource competition, physiology, morphology, and the toxicity of the species, we further investigate how the functional composition and diversity of phytoplankton communities of emerging clusters have changed over time. We hypothesize that environmental filtering over four decades has led to niche separations following the species sorting paradigm in metacommunity ecology (Leibold *et al*. 2004), restructuring community compositions, and resulting in high prevalence of tolerant species and low prevalence among remaining species within genera, i.e. taxonomic dispersion. Exploring the relationship between regions of common community profiles and diversity metrics, at taxonomic and functional levels, enables us to understand how environmental change influences aspects of phytoplankton biodiversity and function on a macrosystem scale (*sensu* Heffernan *et al*. 2014).

## MATERIAL AND METHODS

### Study area and data

We used the Finnish national phytoplankton monitory database maintained by the Finnish Environment Institute (SYKE) that comprises nationwide phytoplankton community data from lake surface water samples taken during the summer months (July, August) between 1977 and 2017 (Open data portal http://www.syke.fi/en-US/Open_information). All phytoplankton samples were preserved with acid Lugol’s solution and analysed using the standard Utermöhl technique (CEN 2006). To avoid computational constraints and maintain model capacity, we selected those species that contributed to the cumulative upper 95% of the total biomass during the four decades of monitoring, resulting in a total of 165 species belonging to 13 classes, and located in 853 lakes belonging to 666 watersheds in 54 river basins. To account for traits in species responses to environmental covariates, we included six traits characterizing important functional aspects of phytoplankton after Weithoff (2003) and Litchman & Klausmeier (2008), namely, cell volume, nitrogen fixation, demand for silica, motility, chain forming ability, and toxicity (Table 1).

**Table 1:** Included phytoplankton traits and presumed reflective functional relevance.

| Trait | Trait category | Functional relationship |
| --- | --- | --- |
| Cell volume | Continuous [$\mu\text{m}^3$] | According to allometric theory, serves as determinant of physiological activities, such as growth and nutrient uptake. |
| Nitrogen fixation | Categorical (yes/no) | The ability of nitrogen fixation provides competitive advantages under nitrogen limitation. |
| Demand for silica | Categorical (yes/no) | Silica increases the specific weight leading to higher sedimentation rates. Besides diatoms that need silica for their frustules, other orders require it for statospores, bristles and scales. |
| Motility | Categorical (yes/no) | Mobile organisms can migrate into favourable patches and counteract sedimentation. Motility also effects nutrient deficiency as the movement of cells minimises the hydrate envelope and, thus, the diffusive boundary layer for nutrients around the cells. |
| Chain forming | Categorical (yes/no) | Larger cell sizes generally experience lower grazing pressure. Hence, the formation of chains, which can be induced by biotic and abiotic factors, may result in reduced mortality due to grazing pressure (Pančić & Kiørboe 2018). |
| Toxicity | Categorical (yes/no) | Toxic phytoplankton species may develop harmful algae blooms (HAB). Such blooms are not only reflecting the impact on water quality, but on species diversity, community structure and ecosystem functioning. Most HABs lead to a significantly decreased diversity and an impairment of many ecosystem functions (Litchman *et al.* 2010). |

In cases where there were no species-specific trait data available, we applied the values of the closest taxonomic level for categorical traits and the mean values of the family level for the continuous cell volume (Weigel *et al*. 2016). We subsequently build a phylogenetic tree of the included species based on their taxonomic structure including class, order, family, genus and species, assuming a branch length of 1 between each node since true phylogenetic information of species was not available.

Related to the long-term nature of the data, spanning four decades, there have been taxonomic changes of species identities stemming from improved identification protocols, keys, and variable identification skills and efforts of phytoplankton analysts. To account for these changes over time, potentially affecting inference on emergence of novel community compositions, we ran a conservative sensitivity analysis including only those species identities that have been consistently classified since the onset of the monitoring campaign. Thus, in cases where e.g., a previously identified species *Aa* was split into multiple species *Aa, Ab, Ac, …* over the course of the study time frame, we converged all subsequent emerging species *Ab, Ac*, … back to the original species, *Aa*, resulting in a total of 133 species instead of 165 (see supplementary material S1 and Table S1 for details).

### Statistical analysis

We analysed the data with hierarchical modelling of species communities (HMSC, Ovaskainen et al., 2017; Ovaskainen & Abrego, 2020). HMSC belongs to the class of joint species distribution models (Warton *et al*. 2015), including a hierarchical layer for how species responses to environmental covariates depend on species traits and phylogenetic relationships (Abrego *et al*. 2017). Here we utilize spatially structured latent variables which were originally proposed by Ovaskainen *et al*. (2016) and later expanded to big spatial data (Tikhonov *et al*. 2020a). For the analyses, we used phytoplankton community data comprising of presence-absence of 165 species surveyed at 1057 sampling locations in 853 lakes across Finland over 40 years from 1977 to 2017. As sampling unit, we used the individual samples taken at each location. Since we modelled presence-absence data, we applied a probit regression model.

As fixed effects we included a suit of environmental predictors which are known to influence phytoplankton communities ranging from physico-chemical water covariates to aspects of lake topography and land use (Table 2). We included water temperature, as well as nutrient concentrations, specifically total phosphorous [μgl^-1^] and the ratio of total nitrogen to total phosphorus and water colour. Covariates related to lake topography were lake-specific surface area (km^2^), mean depth (m) and retention days (the residence time of water in a lake). As proxies for land use extent around lakes, we included the proportions of urban area, agriculture fields and forests in the lakes’ catchment, considering mineral and peat soil types of forests separately, due to their different properties in influencing runoff (Röman *et al*. 2018). To account for temporal trends not described with included covariates, we also added year as linear fixed effect. Since we assumed that species niches may include their optimum at intermediate values of physico-chemical variables, we implemented these covariates as quadratic response functions, while the remaining covariates were assumed to have a linear relationship. To account for the spatial nature of the study design we included spatially explicit random effects at the level of sample site, implemented through the predictive Gaussian process for big spatial data (Tikhonov *et al*. 2020a). To investigate the scale dependent aspects of explained variation as well as species co-occurrence patterns, we structured hierarchical random effects of site, nested in watershed, nested in river basin. To further account for the temporal stochasticity of the data we also included year as random-level effect.

**Table 2:** Included environmental covariates and their relevance for structuring phytoplankton communities

| Covariate group | Covariate | Unit | Relevance |
| --- | --- | --- | --- |
| physico-chemical | water temperature | °C | Temperature directly affects metabolic and growth rates (Padfield *et al.* 2016) and bloom dynamics (Peeters *et al.* 2007). |
|  | total phosphorous (PTOT) concentration | [ugl<sup>-1</sup>] | Phosphorus is a key component determining ecological quality and community structure in lakes (Schindler 1977; Vitousek & Howarth 1991), being a major driver of phytoplankton biomass (Phillips *et al.* 2008) and decreasing biodiversity and water clarity (Carpenter *et al.* 1998; Jeppesen *et al.* 2009). |
|  | total nitrogen to phosphorus ratio (N:P) | unitless ratio | Proxy for nutrient availability. The N:P ratio describes one major factor regulating the dominance of planktonic communities by blue-green or green microorganisms, with a decreasing N:P ratio favouring cyanobacterial blooming (Levich 1996). |
|  | water colour | mg Pt l<sup>-1</sup> | Acts as measure for humic substances, dissolved organic matter and light attenuation (Håkanson 1993; Nürnberg 1998). |
| lake topography | lake area | km<sup>2</sup> | Lakes can be considered as islands (Dodson 1992) with a terrestrial surrounding following the species area theory. Nutrient enrichment effects and phytoplankton diversity are dependent on ecosystem size (Baho *et al.* 2017). |
|  | mean lake depth | m | Lake depth integrates many other habitat features such as water layer mixing, extant of the photic zone and nutrient distribution and is known to impact phytoplankton functional diversity (Longhi & Beisner 2010). |
|  | retention days | days | Retention time influences many aspects of phytoplankton populations (Elliott *et al.* 2009). For example, the rate of water movement through a lake may contribute to the loss and gains of nutrients, organic matter and turbidity (Søballe & Kimmel 1987). |
| land use | urban (or constructed) area | % of catchment area | Catchment characteristics reflect the degree of anthropogenic pressures and sources of nutrients and runoff (Röman *et al.* 2018). |
|  | area of agricultural fields | % of catchment area |  |
|  | mineral forest area | % of catchment area |  |
|  | peat forest area | % of catchment area |  |

HMSC involves a hierarchical structure examining how species responses to environmental covariates depend on species traits and phylogenetic relationships. Hence, we included traits (see Table 1) and the described phylogenetic relationship of the species as a taxonomic tree in our model.

We fitted the HMSC model with the R package Hmsc (Tikhonov et al., 2020b) assuming the default prior distribution (see Chapter 8 of Ovaskainen & Abrego 2020). We sampled the posterior distribution with four Markov chain Monte Carlo (MCMC) chains, each of which was run for 37 500 iterations, of which 12 500 were removed as burn in. The chains were thinned by 100 to yield 250 posterior samples per chain, resulting in 1000 posterior samples in total. We assessed MCMC convergence by examining the potential scale reduction factors (Gelman & Rubin 1992) of the model parameters.

We examined the explanatory power of our model through species specific AUC (Pearce & Ferrier 2000) and Tjur’s R^2^ (Tjur 2009) values, both of which measure how well the model was capable in discriminating between presences and absences. To quantify the drivers of community structure, we then followed Ovaskainen et al. (2017) to partition the explained variation among the fixed and random effects. To illustrate the changes in community composition over space and time, we calculate regions of common community profile based on the predicted species occurrence matrix at the sample level. We used the the *NbClust* package (Charrad *et al*. 2014) to determine an optimal clustering scheme. *NbClust* delivers a framework for determining the best number of clusters, providing 30 indices, and proposing the best clustering scheme obtained from the different methods and results. To highlight relationships of functional diversity and species richness of the emerging clusters, we calculated the functional dispersion (FDis) using the FD package (Laliberté *et al*. 2014) based on predicted community compositions. All analysis was performed in R version 3.6.3 (R Development Core Team 2020).

## RESULTS

### Model convergence and fit

The MCMC convergence was satisfactory. The potential scale reduction factors for the beta-parameters (responses of species to environmental covariates *sensu* Ovaskainen et al. 2017) were on average 1.03 (mean upper C.I. = 1.06). Our model showed a good fit to the data with mean AUC value of 0.88 and mean Tjur’s R^2^ value of 0.23 (see Fig. S1 for details).

### Explained variation in species occurrences

We observed strong spatio-temporal structuring at the species and community levels. About half of the explained variation was attributed to the spatially structured random effects of site ID (18.6.%), watershed (10.8%) and river basin (7%), as well of the temporal random effect of the year (17.8%). Among the fixed effects, the total phosphorus explained most (13.8%) followed by the linear effect of year (10.7%). As none of the remaining fixed effects explained substantial amounts of variation by itself (i.e., explained individually < 5% variation, see supplement Fig. S2), we examined the proportions of variation explained by covariate groups i.e., physico-chemical water parameters (temperature, total N:P, total phosphorus, water colour), lake topography (lake area, mean depth, retention days) and land use (human area, agricultural field, forest type). The physico-chemical water characteristics contributed most to the partitioned variation with 18%, followed by land use with 9.4%, and lake topography contributed with 7%. The species traits explained on average 6.7% of the among-species variation in species responses to the fixed effects, with the strongest effect (21% of explained variation) related to species responses to lake area (21%) (Fig. S3).

### Species responses to covariates and phylogenetic niche conservatism

The species responses to environmental covariates showed phylogenetic structuring with strong statistical support for the hypothesis of niche conservatism, reflected in the posterior mean of the phylogenetic correlation parameter ρ being 0.84 (with a 95% credible interval from 0.76 to 0.91). Hence, closely related species tend to respond similarly to the covariates included in the model (Fig. 1). Accounting only for species occurrence responses that were positive or negative with at least 95% posterior probability, the linear effect of year influenced most of the species’ occurrences. Temperature affected one fifth of the species occurrences, with positive linear and negative quadratic response. More than half of all species occurrences were positively affected by the linear effect of total phosphorus concentrations, while the second order response was mainly negative. Total phosphorus constituted the strongest physico-chemical predictor for the metacommunity composition. Considering aspects of lake topography, almost two thirds of the species showed mainly positive associations with total lake area, while retention days and lake depth had negative effect for occurrences of half of the species. Statistically supported responses to the size of urban areas and agricultural fields were mainly positive. When investigating the impact of different forest soil types in the catchment area, we found mineral forest soils to affect phytoplankton occurrences positively while peat forest soils had a negative impact (Fig.1).

**Figure 1:**
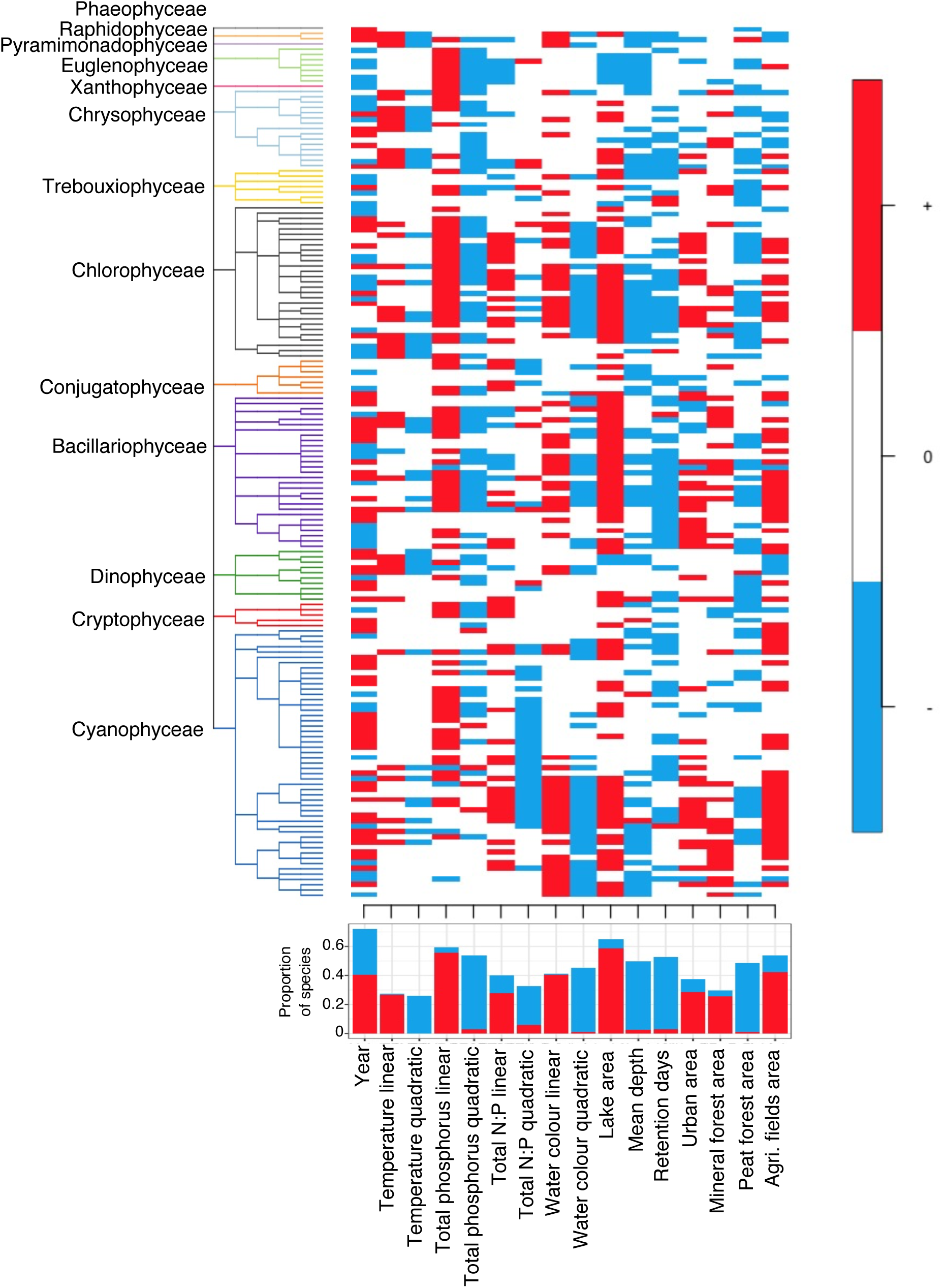
Phylogenetically structured responses of species occurrences to the covariates. Response that are positive in at least 95% posterior probability are shown by red, responses that are negative with at least 95% posterior probability are shown in blue. Responses that did not gain strong statistical support are shown in white. Bar plot below summarizes the proportion of species responses being positive or negative for each covariate.

### Spatial scales of species variation

We found species co-occurrences to display highly scale-dependent patters. At the local level of sampling location, our results show a large proportion of species positively co-occurring (Fig 2). When increasing the spatial scale to the level of watershed, there is a clear decrease in species co-occurrences with high statistical support as well as an increase in species being negatively associated, a pattern intensifying when increasing the spatial scale to river basin. At the level of year, associations of species were clearly indicating that the stochastic effect of time resulted in strong negative and positive residual associations of species, representative of temporal community change.

**Figure 2:**
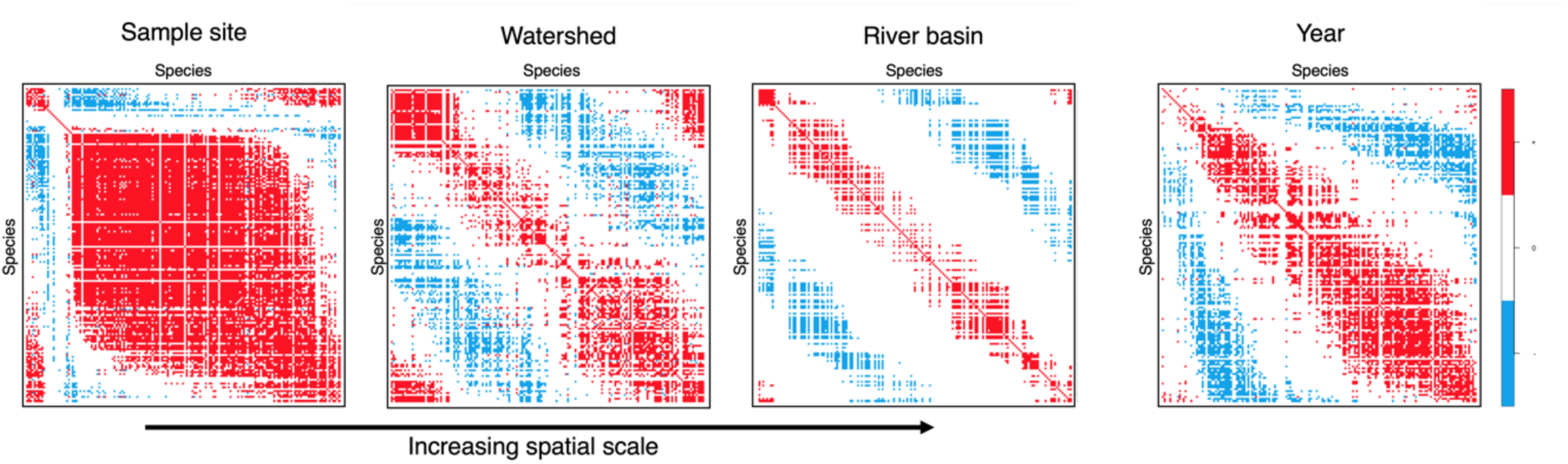
Residual association among species at spatial and temporal scales. Pairs of species illustrated by red and blue show positive and negative associations, respectively, with statistical support of at least 95% posterior probability.

### Spatio-temporal change in taxonomical composition

Above, we have reported how phytoplankton communities respond to variation in environmental conditions, identifying the total amount of phosphorus as a key driver of community composition. During the four decades that our data span, lake ecosystems have undergone major environmental changes due to anthropogenic activities, resulting in a major change in community composition (Fig. 3a), illustrated by changing regions of common profile (RCP). Four emerging RCP-clusters resulted from clustering algorithm and are highlighted based on the predicted community composition at the level of sample ID. While RCP cluster 1 was dominant during the mid 70-80s throughout Finland, a complete restructuring took place resulting in RCP 1 disappearing after the mid 90s and a novel RCP cluster 2 and previously scarce cluster 3 becoming dominant with RCP cluster 4 also becoming more common and expanding from only a few lakes in southern Finland northward (Fig. 3a).

**Figure 3:**
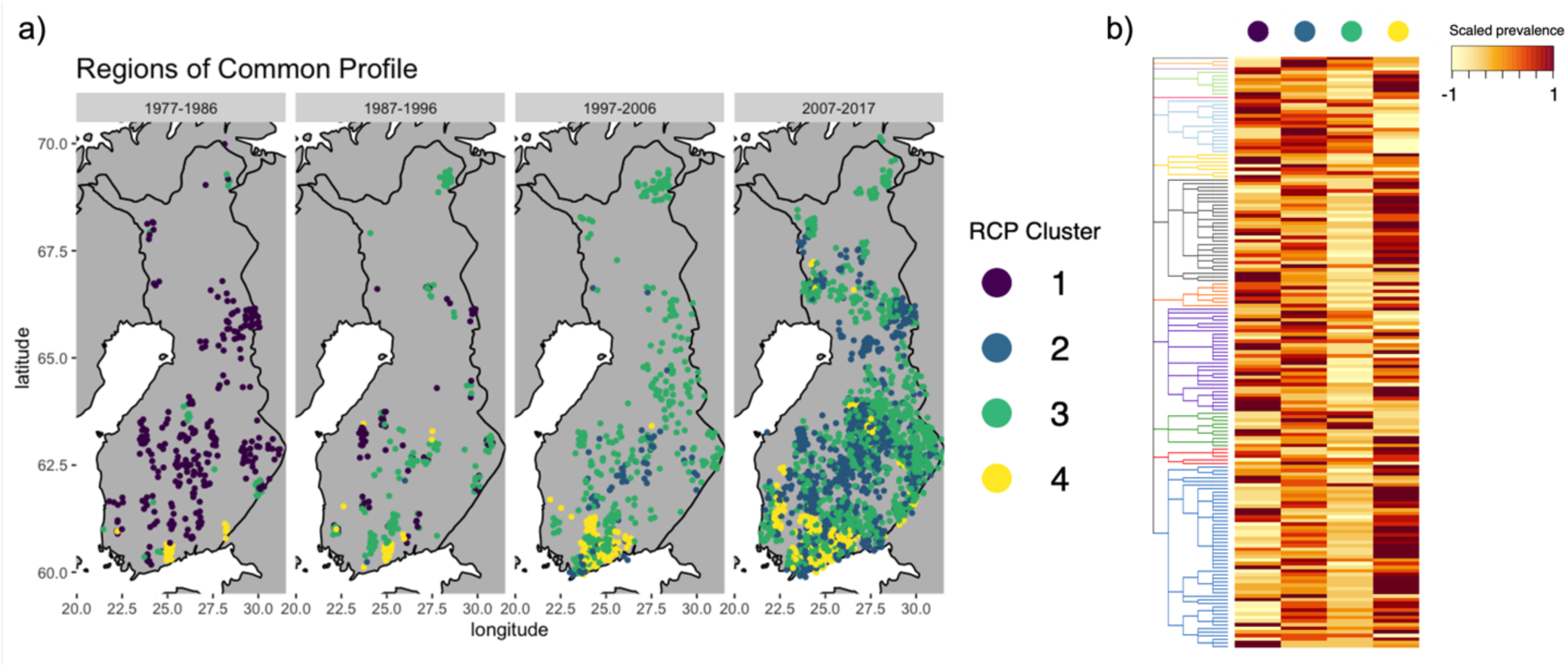
a) Regions of common profile based on optimal clustering of predicted species compositions in lakes. Data are displayed in aggregations of 10 years to highlight the temporal aspect of change. To avoid overlapping data points, the data are plotted with jitter of 0.2 degree for latitude and longitude, b) scaled prevalence of species in each RCP, following the phylogenetic structure of Fig 1.

We looked at the characteristics of the different clusters in terms of species prevalence, species richness, community weighted traits, and the relationship between species richness and functional diversity. Each RCP was associated with different prevalence patterns of species with, in part, strong phylogenetic structuring (Fig. 3b). While RCP 1 showed high prevalence of few dominant species within several taxonomic classes, RCP 4 is characterized by high prevalence of all species comprised in the classes of Cyanophyceae (Cyanobacteria) and Chlorophyceae (green algae) as well as Euglenophyceae and Cryptophyceae (both flagellated). RCP 2 shows no clear phylogenetic structuring in turns of prevalence but depicts overall high species prevalence. This contrasts with RCP 3 that shows generally low prevalence of species, with exception of Dinophyceae (dinoflagellates) and Chrysophyceae (golden algae). We found that mean species richness differed among all RCPs but 1 and 4, with RCP 2 displaying highest and RCP3 lowest richness (Fig 4a, S4). The RCPs also differed by their community weighted traits. Comparing RCPs 1, 2 and 3 we find decreasing proportions of their toxicity (Fig. 4b), silica requirements (with non-significant difference between RCP2 and 3) (Fig 4c, S4), chain forming (Fig. 4d) and nitrogen fixating species (Fig 4e) as well as a decrease in cell volume (Fig. 4g). In contrast, we found an increase in motile species from RCP 1-3 with the latter showing the highest degree of motility (Fig 4f). RCP 4 differed substantially from the other RCPs with its toxicity, chain forming and nitrogen fixation traits being significantly higher and the requirement for silica, motility and cell volume being significantly lower than those in RCPs 1-3 (Fig. 4b-g, S4).

**Figure 4:**
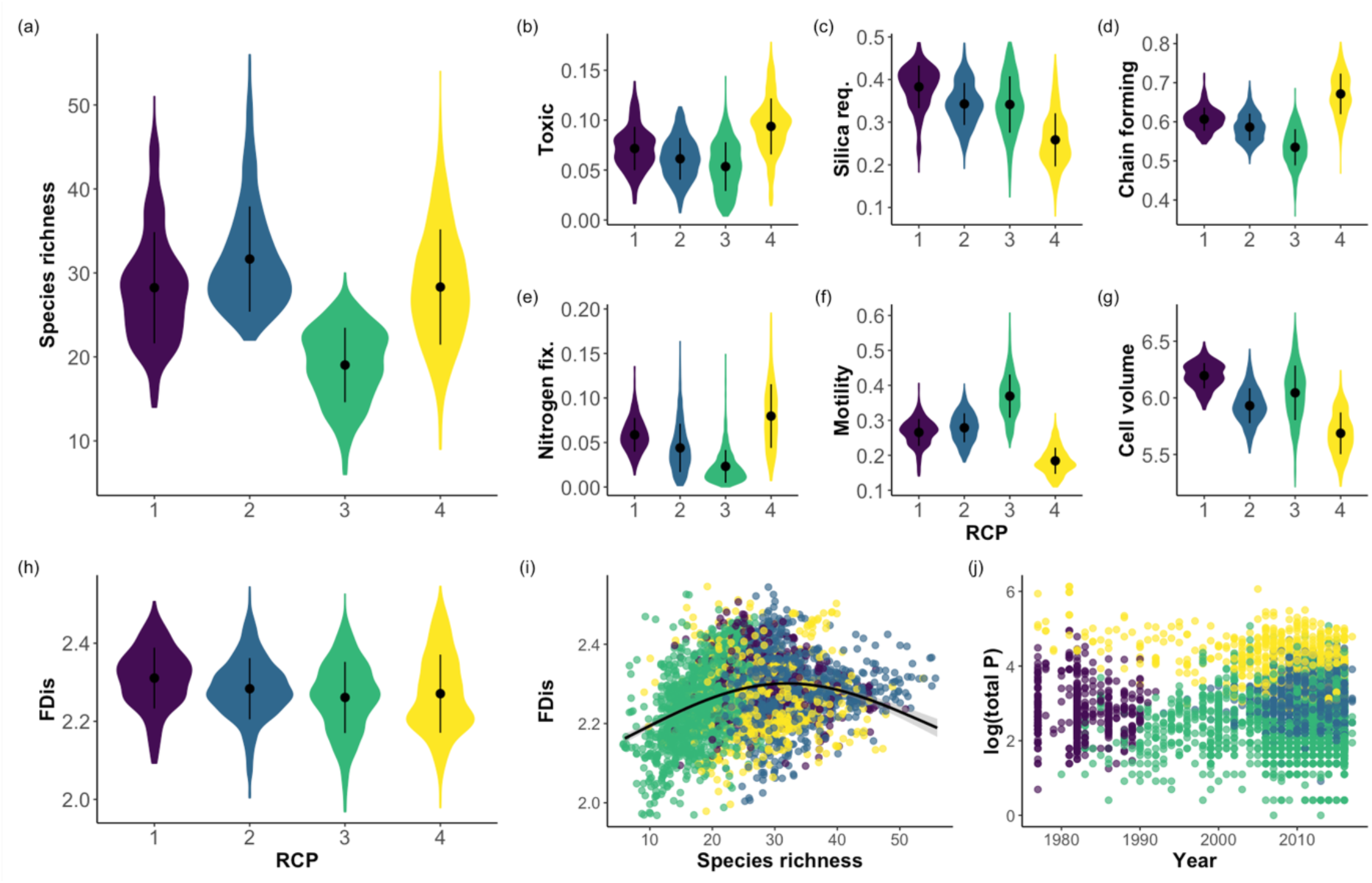
Illustration of community characteristics at the four Regions of Common Profile (RCP). (a) Species richness (b-g) community weighted trait values, (h) Functional diversity displayed as Functional Dispersion FDis, (i) relationship between species richness and functional diversity (FDis), trendline highlights the relationship based on a generalized additive model including all data with 2 knots, (j) total phosphorus concentrations at RCP over time. Colours of RCPs are the same as in Fig. 3.

The functional diversity, measured as functional dispersion (FDis), was highest in RCP 1 and showed more variation in RCPs 2-4, but with overall small changes between the RCPs (Fig. 4h, S4), despite the differences in species richness. On average, maximum FDis values were observed in communities with medium levels of species richness of ca. 30 species, after which higher species richness values resulted in lower functional diversity (Fig 4i), suggesting functional redundancy.

Linking the RCPs to environmental characteristics, we find that RCP 4 is mainly associated with high total phosphorus concentrations (PTOT) over the entire study period, while RCP 3 is more prevalent in the mid and especially lower ranges of PTOT (Fig. 4j). RCPs 1 and 2, representing the originally dominating and novel cluster, respectively, are most prevalent in between highest (RCP 4) and lowest (RCP 3) phosphorus contractions (Fig. 4j).

Despite some changes in taxonomical identification procedures that affected 32 of the initial 165 species (mainly after the year 2000) our sensitivity analysis only considering the previous species complex of 133 species shows the same clustering results (Fig. S5), supporting the robustness of our model and the strong temporal signal of community change we detected.

## DISCUSSION

The concept of lakes as meta-systems helps to structure the large-scale impacts of environmental change on biodiversity and ecosystem functions (Heino *et al*. 2020). Here, we have shown that a nationwide restructuring of lake phytoplankton communities took place in Finland over the past four decades. The historically most dominant and widespread community profile has vanished over time, now being replaced by two previously rarer and one novel community profile (Fig. 3a). The emerging distribution of different regions of common profiles (RCP) are a result from deviating community compositions, stemming from abiotic and biotic filtering on species levels, in a macrosystem spatio-temporal context. We show that the interplay between environmental covariates and species occurrences, i.e., species niches, display a robust phylogenetic structure. Within these structured responses, species prevalence shifted from high taxonomic dispersion, i.e., only few highly prevalent (dominant) species per class, to low taxonomic dispersion, i.e., nearly all species with similar prevalence within classes, comparing the previously dominant with the more recent community profiles (Fig. 3b). This contrasts with our assumption that over time, taxonomic dispersion, i.e. the prevalence of only a few tolerant species within genera and families, would increase under strong environmental filtering (Passy *et al*. 2017). Instead, we observed lower taxonomic dispersion with overall more homogeneous, but community type-specific species prevalence, that could be explained by the strong phylogenetic niche conservatisms. This phylogenetic signal implies phylogenetically correlated response traits, that ultimately determine the species niche.

In our work, the included traits reflecting aspects of resource competition, physiology, morphology, and toxicity, did not explain major proportions of the variation among the species in their responses to the environmental covariates. However, distinct combinations of traits among species may support different responses to changes in the environment as a result of environmental filtering under altered niche spaces (Litchman *et al*. 2012; Mouillot *et al*. 2013). The differences in community weighted mean traits among the four RCPs indeed suggest altered community characteristics, and hence, presumably also functioning. Especially the communities in RCP 4 differed from the rest (Fig 3b-j). RCP 4 was exclusively found in environments with high total phosphorus concentrations (Fig 3j). In these conditions nitrogen fixation ability is beneficial for species as indicated by the highest community weighted mean of nitrogen fixation (nfix trait) in this RCP, potentially giving the species complex competitive advantage under nitrogen limitation. Furthermore, RCP 4 showed lowest community weighted means for silica requirement, motility, and cell volume, while displaying highest proportions of toxic and chain forming species. All these characteristics point to a community dominated by Cyanobacteria (Cyanophyceae), a functional group that may cause harmful algae blooms and is, due to its toxicity and morphological traits, well known for its lower nutritional quality and accessibility as food resource for zooplankton and other grazers, able to decouple trophic links and alter food web structure (Elser & Goldman 1991; Wilson *et al*. 2006). However, compared to the historically dominant RCP 1, the emerging and currently dominant RCPs 2-4 also show on average smaller cell volumes (Fig. 3g). Smaller phytoplankton size structures have been linked to increased temperatures due to climate change (Winder *et al*. 2009; Mousing *et al*. 2014; Zohary *et al*. 2021), which may explain why the more recent community types are smaller compared the historically dominant type of the late 70s and 80s. From an ecosystem functioning perspective, this may become problematic as smaller phytoplankton is commonly favoured by smaller zooplankton predators, altering the food chain length and resulting in less favourable resources also for higher trophic levels (Litchman *et al*. 2015).

In our work, we highlight both abiotic and biotic filtering processes of phytoplankton communities by modelling the species-specific environmental drivers through the implemented fixed effects, and aspects of biotic filtering through the random effect part in our model. While physico-chemical water characteristics jointly contributed most to the explained variation at the community level, the largest explained variation is related to spatial and temporal random effects. These random effects can be linked to biotic filtering, i.e. how the ecological interactions among species influence their occurrences, particularly their co-occurrences (Fig. 2), but also to missing random level specific covariates (Ovaskainen & Abrego 2020). Considering the spatial scales from smallest (sample location) to largest (river basin), we found that the hierarchical spatial effects are decreasing in their importance of explaining species occurrences with increasing scale. This is in line with other findings (Heino *et al*. 2015; Ovaskainen *et al*. 2019) suggesting that connectivity and biotic filtering decrease towards larger scales. Thus, with increasing scales, local abiotic drivers are progressively governed by regional drivers and the subsequent cross-scale interactions among these drivers increase the macroscale complexity (Soranno *et al*. 2014).

The relatively strong temporal signal, reflected here in the linear effect of year (fixed effect) also suggests that there may be a set of important unmeasured environmental parameters that have not been included in our study, such as more detailed inorganic nutrient fractions that have been shown to explain phytoplankton community properties partly better than only including total concentrations (Ptacnik *et al*. 2010; Trommer *et al*. 2020). Such detailed data is unfortunately scarce especially over broad spatio-temporal scales. However, we find strong associations between the included environmental predictors and species occurrences that exceed the explained variation of the linear temporal signal. While temperature showed overall strong associations with relatively few species, nutrient concentrations showed most frequent associations with species occurrences. When also considering aspects of land use in the catchment of the lakes, we found generally uniform responses among species, showing mainly positive associations with urban, mineral forest and agricultural areas, all being linked to higher nutrient inputs. However, peat forest area showed virtually only negative associations (Fig. 1), which may be linked to more dissolved organic carbon in the runoff, and the “browning” of lakes, hindering phytoplankton production through light absorption and shading (Klug 2002; Thrane *et al*. 2014). Our results support this relationship further by displaying strong relationships with water colour as proxy for water clarity. Dissolved organic carbon levels are increasing due to climate change and land-use (peat mining and forestry), an issue that has been neglected in water management until recently (Kritzberg *et al*. 2020). The lake water colour also affects the critical phosphorus thresholds that triggers the growth of bloom-forming cyanobacteria (Vuorio *et al*. 2020) and decreases the nutritional quality of fish for human consumers (Taipale *et al*. 2016).

When considering the historically vastly dominating community type of RCP 1 as a baseline, we show that environmental filtering and species sorting have resulted in a macrosystem-wide restructuring of Finnish lake phytoplankton communities over the past four decades. This may have potentially wide-reaching implications for lake ecosystem functioning, both for species interactions, e.g., food web structure, as well as for human well-being, e.g. proportion of present toxic species. When considering lakes as metacommunity hubs in an interconnected waterscape, we show that the assembly mechanisms of phytoplankton communities are strongly structured by spatial and temporal dynamics, leading to novel community types under progressive environmental change. Since the establishment of such restructuring happens over several decades and can only be showcased on large spatial scales, it will be important to consider how different trajectories of environmental change may intensify observed developments in community composition and how these may be reflected in water management under the European Water Framework Directive that depend on community compositions and evidence-driven decision-making.

## ACKNOWLEDGEMENTS

The study was funded by the Strategic Research Council of Academy of Finland (Project 312650 BlueAdapt) and supported by Academyof Finland grants 309581 (OO) and 311229 (KV) and the European Research Council (ERC) under the European Union’s Horizon 2020 research and innovation programme; grant agreement No 856506; ERC-synergy project LIFEPLAN) (OO). We thank Elina Kaarlejärvi for comments on an earlier version of the manuscript.

## REFERENCES

Abrego, N., Norberg, A. & Ovaskainen, O. (2017). Measuring and predicting the influence of traits on the assembly processes of wood-inhabiting fungi. J. Ecol., 105, 1070–1081.

Baattrup-Pedersen, A., Larsen, S.E., Rasmussen, J.J. & Riis, T. (2019). The future of European water management: Demonstration of a new WFD compliant framework to support sustainable management under multiple stress. Sci. Total Environ., 654, 53–59.

Baho, D.L., Drakare, S., Johnson, R.K., Allen, C.R. & Angeler, D.G. (2017). Is the impact of eutrophication on phytoplankton diversity dependent on lake volume/ecosystem size? J. Limnol., 76, 199–210.

Cadotte, M.W., Arnillas, C.A., Livingstone, S.W. & Yasui, S.L.E. (2015). Predicting communities from functional traits. Trends Ecol. Evol., 30, 510–511.

Carpenter, S.R., Caraco, N.F., Correll, D.L., Howarth, R.W., Sharpley, A.N. & Smith, V.H. (1998). Nonpoint pollution of surface waters with phosphorus and nitrogen. Ecol. Appl., 8, 559–568.

Carvalho, L., Mackay, E.B., Cardoso, A.C., Baattrup-Pedersen, A., Birk, S., Blackstock, K.L., et al. (2019). Protecting and restoring Europe’s waters: An analysis of the future development needs of the Water Framework Directive. Sci. Total Environ., 658, 1228–1238.

CEN. (2006). EN 15204 − Water quality − Guidance standard on the enumeration of phytoplankton using inverted microscopy (Utermöhl technique)., 1–39.

Charrad, M., Ghazzali, N., Boiteau, V. & Niknafs, A. (2014). Nbclust: An R package for determining the relevant number of clusters in a data set. J. Stat. Softw., 61, 1–36.

Chase, J.M. & Leibold, M.A. (2002). Spatial scale dictates the productivity-biodiversity relationship. Nature, 416, 427–430.

Chase, J.M., McGill, B.J., Thompson, P.L., Antão, L.H., Bates, A.E., Blowes, S.A., et al. (2019). Species richness change across spatial scales. Oikos, 1079–1091.

Dodson, S. (1992). Predicting crustacean zooplankton species richness. Limnol. Oceanogr., 37, 848–856.

Dudgeon, D., Arthington, A.H., Gessner, M.O., Kawabata, Z.I., Knowler, D.J., Lévêque, C., et al. (2006). Freshwater biodiversity: Importance, threats, status and conservation challenges. Biol. Rev. Camb. Philos. Soc., 81, 163–182.

Edwards, K.F., Litchman, E. & Klausmeier, C.A. (2013). Functional traits explain phytoplankton responses to environmental gradients across lakes of the United States. Ecology, 94, 1626–1635.

Elliott, J.A., Jones, I.D. & Page, T. (2009). The importance of nutrient source in determining the influence of retention time on phytoplankton: An explorative modelling study of a naturally well-flushed lake. Hydrobiologia, 627, 129–142.

Elser, J.J. & Goldman, C.R. (1991). Zooplankton effects on phytoplankton in lakes of contrasting trophic status. Limnol. Oceanogr., 36, 64–90.

European Comission. (2000). Directive 2000/60/EC of the European Parliament and of the Council of 23 October 2000 establishing a framework for community action in the field of water policy. Official. Off. J. Eur. Communities, L 327/1-L327/72.

Falkenmark, M. (2003). Freshwater as shared between society and ecosystems: From divided approaches to integrated challenges. Philos. Trans. R. Soc. B Biol. Sci., 358, 2037–2049.

Gagic, V., Bartomeus, I., Jonsson, T., Taylor, A., Winqvist, C., Fischer, C., et al. (2015). Functional identity and diversity of animals predict ecosystem functioning better than species-based indices. Proc. R. Soc. B, 282, 20142620.

Gelman, A. & Rubin, D.B. (1992). Inference from Iterative Simulation Using Multiple Sequences. Stat. Sci., 2, 45–52.

Grizzetti, B., Liquete, C., Pistocchi, A., Vigiak, O., Zulian, G., Bouraoui, F., et al. (2019). Relationship between ecological condition and ecosystem services in European rivers, lakes and coastal waters. Sci. Total Environ., 671, 452–465.

Håkanson, L. (1993). A Model to Predict Lake Water Colour. Int. Rev. der gesamten Hydrobiol. und Hydrogr., 78, 107–137.

Hardin, G. (1960). The competitive exclusion principle. Science (80-.)., 131, 1292–1297.

Harvey, P.H. & Purvis, A. (1991). Comparative methods for explaining adaptations. Nature, 351, 619–624.

Heffernan, J.B., Soranno, P.A., Angilletta, M.J., Buckley, L.B., Gruner, D.S., Keitt, T.H., et al. (2014). Macrosystems ecology: Understanding ecological patterns and processes at continental scales. Front. Ecol. Environ., 12, 5–14.

Heino, J., Alahuhta, J., Bini, L.M., Cai, Y., Heiskanen, A., Hellsten, S., et al. (2020). Lakes in the era of global change: moving beyond single-lake thinking in maintaining biodiversity and ecosystem services. Biol. Rev., 4, brv.12647.

Heino, J., Melo, A.S., Siqueira, T., Soininen, J., Valanko, S. & Bini, L.M. (2015). Metacommunity organisation, spatial extent and dispersal in aquatic systems: Patterns, processes and prospects. Freshw. Biol., 60, 845–869.

Jeppesen, E., Kronvang, B., Meerhoff, M., Søndergaard, M., Hansen, K.M., Andersen, H.E., et al. (2009). Climate Change Effects on Runoff, Catchment Phosphorus Loading and Lake Ecological State, and Potential Adaptations. J. Environ. Qual., 38, 1930–1941.

Keddy, P.A. (1992). Assembly and response rules: two goals for predictive community ecology. J. Veg. Sci., 3, 157–164.

Klug, J.L. (2002). Positive and negative effects of allochthonous dissolved organic matter and inorganic nutrients on phytoplankton growth. Can. J. Fish. Aquat. Sci., 59, 85–95.

Kritzberg, E.S., Hasselquist, E.M., Škerlep, M., Löfgren, S., Olsson, O., Stadmark, J., et al. (2020). Browning of freshwaters: Consequences to ecosystem services, underlying drivers, and potential mitigation measures. Ambio, 49, 375–390.

Laliberté, E., Legendre, P. & Shipley, B. (2014). FD: measuring functional diversity from multiple traits, and other tools for functional ecology. R package version 1.0-12.

Leibold, M.A., Holyoak, M., Mouquet, N., Amarasekare, P., Chase, J.M., Hoopes, M.F., et al. (2004). The metacommunity concept: A framework for multi-scale community ecology. Ecol. Lett., 7, 601–613.

Lepistö, L., Holopainen, A.L. & Vuoristo, H. (2004). Type-specific and indicator taxa of phytoplankton as a quality criterion for assessing the ecological status of Finnish boreal lakes. Limnologica, 34, 236–248.

Levich, A.P. (1996). The role of nitrogen-phosphorus ratio in selecting for dominance of phytoplankton by cyanobacteria or green algae and its application to reservoir management. J. Aquat. Ecosyst. Stress Recover., 5, 55–61.

Litchman, E., Edwards, K.F., Klausmeier, C.A. & Thomas, M.K. (2012). Phytoplankton niches, traits and eco-evolutionary responses to global environmental change. Mar. Ecol. Prog. Ser., 470, 235–248.

Litchman, E. & Klausmeier, C.A. (2008). Trait-based community ecology of phytoplankton. Annu. Rev. Ecol. Evol. Syst., 39, 615–639.

Litchman, E., de Tezanos Pinto, P., Edwards, K.F., Klausmeier, C.A., Kremer, C.T. & Thomas, M.K. (2015). Global biogeochemical impacts of phytoplankton: A trait-based perspective. J. Ecol., 103, 1384–1396.

Litchman, E., de Tezanos Pinto, P., Klausmeier, C.A., Thomas, M.K. & Yoshiyama, K. (2010). Linking traits to species diversity and community structure in phytoplankton. Hydrobiologia, 653, 15–28.

Longhi, M.L. & Beisner, B.E. (2010). Patterns in taxonomic and functional diversity of lake phytoplankton. Freshw. Biol., 55, 1349–1366.

Mammides, C. (2020). A global assessment of the human pressure on the world’s lakes. Glob. Environ. Chang., 63, 102084.

Mayfield, M.M. & Levine, J.M. (2010). Opposing effects of competitive exclusion on the phylogenetic structure of communities. Ecol. Lett., 13, 1085–1093.

McGill, B.J., Enquist, B.J., Weiher, E. & Westoby, M. (2006). Rebuilding community ecology from functional traits. Trends Ecol. Evol. (Personal Ed., 21, 178–85.

Mouillot, D., Graham, N. a J., Villéger, S., Mason, N.W.H. & Bellwood, D.R. (2013). A functional approach reveals community responses to disturbances. Trends Ecol. Evol., 28, 167–77.

Mousing, E.A., Ellegaard, M. & Richardson, K. (2014). Global patterns in phytoplankton community size Structure-evidence for a direct temperature effect. Mar. Ecol. Prog. Ser., 497, 25–38.

Myers, N. (2003). Biodiversity Hotspots Revisited. Bioscience, 53, 916–917.

Nürnberg, G.K. (1998). Productivity of clear and humic lakes: Nutrients, phytoplankton, bacteria. Hydrobiologia, 382, 151–159.

Ovaskainen, O. & Abrego, N. (2020). Joint Species Distribution Modelling. Cambridge University Press.

Ovaskainen, O., Abrego, N., Halme, P. & Dunson, D. (2016). Using latent variable models to identify large networks of species-to-species associations at different spatial scales. Methods Ecol. Evol., 7, 549–555.

Ovaskainen, O., Tikhonov, G., Dunson, D., Grøtan, V., Engen, S., Sæther, B., et al. (2017). How are species interactions structured in species-rich communities ? A new method for analysing time-series data. Proc. R. Soc. B, 284, 20170768.

Ovaskainen, O., Weigel, B., Potyutko, O. & Buyvolov, Y. (2019). Long-term shifts in water quality show scale-dependent bioindicator responses across Russia – Insights from 40 year-long bioindicator monitoring program. Ecol. Indic., 98, 476–482.

Padfield, D., Yvon-Durocher, G., Buckling, A., Jennings, S. & Yvon-Durocher, G. (2016). Rapid evolution of metabolic traits explains thermal adaptation in phytoplankton. Ecol. Lett., 19, 133–142.

Padisák, J., Borics, G., Grigorszky, I. & Soróczki-Pintér, É. (2006). Use of Phytoplankton Assemblages for Monitoring Ecological Status of Lakes within the Water Framework Directive: The Assemblage Index. Hydrobiologia, 553, 1–14.

Pančić, M. & Kiørboe, T. (2018). Phytoplankton defence mechanisms: traits and trade-offs. Biol. Rev., 93, 1269–1303.

Passy, S.I., Bottin, M., Soininen, J. & Hillebrand, H. (2017). Environmental filtering and taxonomic relatedness underlie the species richness–evenness relationship. Hydrobiologia, 787, 243–253.

Pearce, J. & Ferrier, S. (2000). Evaluating the predictive performance of habitat models developed using logistic regression. Ecol. Modell., 133, 225–245.

Peeters, F., Straile, D., Lorke, A. & Livingstone, D.M. (2007). Earlier onset of the spring phytoplankton bloom in lakes of the temperate zone in a warmer climate. Glob. Chang. Biol., 13, 1898–1909.

Phillips, G., Pietiläinen, O.P., Carvalho, L., Solimini, A., Lyche Solheim, A. & Cardoso, A.C. (2008). Chlorophyll-nutrient relationships of different lake types using a large European dataset. Aquat. Ecol., 42, 213–226.

Ptacnik, R., Andersen, T. & Tamminen, T. (2010). Performance of the Redfield Ratio and a Family of Nutrient Limitation Indicators as Thresholds for Phytoplankton N vs. P Limitation. Ecosystems, 13, 1201–1214.

R Development Core Team. (2020). R: A language and environment for statistical computing. R Foundation for Statistical Computing, Vienna, Austria. http://www.r-project.org.

Röman, E., Ekholm, P., Tattari, S., Koskiaho, J. & Kotamäki, N. (2018). Catchment characteristics predicting nitrogen and phosphorus losses in Finland. River Res. Appl., 34, 397–405.

Schindler, A.D.W. (1977). Evolution of Phosphorus Limitation in Lakes Published by : American Association for the Advancement of Science Stable URL : http://www.jstor.org/stable/1743244 Accessed : 29-05-2016 18 : 58 UTC. Science (80-.)., 195, 260–262.

Søballe, D.M. & Kimmel, B.L. (1987). A Large-Scale Comparison of Factors Influencing Phytoplankton Abundance in Rivers, Lakes, and Impoundments. Ecology, 68, 1943–1954.

Soranno, P.A., Cheruvelil, K.S., Bissell, E.G., Bremigan, M.T., Downing, J.A., Fergus, C.E., et al. (2014). Cross-scale interactions: Quantifying multi-scaled cause-effect relationships in macrosystems. Front. Ecol. Environ., 12, 65–73.

Soranno, P.A., Webster, K.E., Riera, J.L., Kratz, T.K., Baron, J.S., Bukaveckas, P.A., et al. (1999). Spatial variation among lakes within landscapes: Ecological organization along lake chains. Ecosystems, 2, 395–410.

Taipale, S.J., Vuorio, K., Strandberg, U., Kahilainen, K.K., Järvinen, M., Hiltunen, M., et al. (2016). Lake eutrophication and brownification downgrade availability and transfer of essential fatty acids for human consumption. Environ. Int., 96, 156–166.

Thrane, J.E., Hessen, D.O. & Andersen, T. (2014). The Absorption of Light in Lakes: Negative Impact of Dissolved Organic Carbon on Primary Productivity. Ecosystems, 17, 1040–1052.

Tikhonov, G., Duan, L., Abrego, N., Newell, G., White, M., Dunson, D., et al. (2020a). Computationally efficient joint species distribution modeling of big spatial data. Ecology, 101, 1–8.

Tikhonov, G., Opedal, Ø.H., Abrego, N., Lehikoinen, A., Jonge, M.M.J., Oksanen, J., et al. (2020b). Joint species distribution modelling with the r-package Hmsc. Methods Ecol. Evol., 11, 442–447.

Tjur, T. (2009). Coefficients of determination in logistic regression models - A new proposal: The coefficient of discrimination. Am. Stat., 63, 366–372.

Trommer, G., Poxleitner, M. & Stibor, H. (2020). Responses of lake phytoplankton communities to changing inorganic nitrogen supply forms. Aquat. Sci., 82, 1–13.

Vitousek, P.M. & Howarth, R.W. (1991). Nitrogen limitation on land and in the sea: How can it occur? Biogeochemistry, 13, 87–115.

Vuorio, K., Järvinen, M. & Kotamäki, N. (2020). Phosphorus thresholds for bloom-forming cyanobacterial taxa in boreal lakes. Hydrobiologia, 847, 4389–4400.

Warton, D.I., Blanchet, F.G., O’Hara, R.B., Ovaskainen, O., Taskinen, S., Walker, S.C., et al. (2015). So Many Variables: Joint Modeling in Community Ecology. Trends Ecol. Evol., 30, 766–779.

Weigel, B., Blenckner, T. & Bonsdorff, E. (2016). Maintained functional diversity in benthic communities in spite of diverging functional identities. Oikos, 125, 1421–1433.

Weithoff, G. (2003). The concepts of “plant functional types” and “functional diversity” in lake phytoplankton - A new understanding of phytoplankton ecology? Freshw. Biol., 48, 1669–1675.

Williamson, C.E., Dodds, W., Kratz, T.K. & Palmer, M.A. (2008). Lakes and streams as sentinels of environmental change in terrestrial and atmospheric processes. Front. Ecol. Environ., 6, 247–254.

Wilson, A.E., Sarnelle, O. & Tillmanns, A.R. (2006). Effects of cyanobacterial toxicity and morphology on the population growth of freshwater zooplankton: Meta-analyses of laboratory experiments. Limnol. Oceanogr., 51, 1915–1924.

Winder, M., Reuter, J.E. & Schladow, S.G. (2009). Lake warming favours small-sized planktonic diatom species. Proc. R. Soc. B Biol. Sci., 276, 427–435.

Zohary, T., Flaim, G. & Sommer, U. (2021). Temperature and the size of freshwater phytoplankton. Hydrobiologia, 848, 143–155.

