## supplementary material for "Macrosystem community change in lake phytoplankton and its implications for diversity and function"

**S1 Sensitivity analysis to account for improved taxonomical identification**

Due to the broad scale and long-term nature of used community data, comprising four decades, we fitted a separate joint species distribution model, only considering those taxonomic identities that have not changed due to improved identification tools over time. This was to ensure the robustness of our results concerning the clustering and emerging differences in region of common community profiles (RCPs).

Taxonomy is not static and always strives to become more accurate. With international expert exchange and improved identification keys, some species can be identified more accurately today than several decades ago or have simply changed their name in accordance with the latest knowledge of taxonomic classifications. This remains a challenge when working on long-term data sets. The latter, i.e. changed taxonomic names, are easy to fix when curating the data set. However, in cases where identification became more precise and species *Aa* can now be identified into an entirely different species complex, e.g., *Aa, Ab* and *Ac,* this is a bigger challenge. With no means of disentangling this species complex prior to the new classification, we took the revered, and more conservative approach, of regrouping the emerging species back to the initial recorded species identity. In practice this means in cases where e.g., a previously identified species *Aa* has split into multiple species *Aa*, *Ab*, *Ac,…* over the course of the study time frame, we converged all subsequent emerging species *Aa, Ab, Ac*, …back to the original species, *Aa,* resulting in a total of 133 species instead of 165 (Table S1). Most of the changes in species identification happened after 2000. The back transformation to the original species complex is based on expert judgement.

The model structure of the described sensitivity model is the same as for the full model demonstrated in the main manuscript. We also followed an identical procedure, hence sampling the posterior distribution with four Markov chain Monte Carlo (MCMC) chains, each of which was run for 37 500 iterations, of which 12 500 were removed as burn in. The chains were thinned by 100 to yield 250 posterior samples per chain, resulting in 1000 posterior samples in total. Our results show close to identical clustering of RCPs suggesting that the changes in taxonomic classification did not affect the strong changes in RCP clustering over time (Figure S1.)

**Table S1:** List of all 165 included species in the main analysis (left) and list of adjusted species complex for community composition sensitivity analysis (133 entities, right). Highlighted in bold font are species that have changed in identification around the year 2000 and have been regrouped to their previous species identity shown in the right column.

| Full species list | Adjusted species complex |
|---|---|
| *Acanthoceras zachariasii* | *Acanthoceras zachariasii* |
| *Acutodesmus acuminatus* | *Acutodesmus acuminatus* |
| *Anabaena minderi* | *Anabaena minderi* |
| ***Anabaena planctonica*** | ***Dolichospermum planctonicum*** |
| ***Anabaena spiroides*** | ***Dolichospermum spiroides*** |
| ***Anathece bachmannii*** | ***Anathece clathrata cx*** |
| ***Anathece clathrata*** | ***Anathece clathrata cx*** |
| ***Anathece minutissima*** | ***Anathece clathrata cx*** |
| *Ankistrodesmus fusiformis* | *Ankistrodesmus fusiformis* |
| ***Aphanizomenon flexuosum*** | ***Aphanizomenon flosaquae cx*** |
| ***Aphanizomenon flosaquae*** | ***Aphanizomenon flosaquae cx*** |
| ***Aphanizomenon gracile*** | ***Aphanizomenon gracile cx*** |
| ***Aphanizomenon klebahnii*** | ***Aphanizomenon flosaquae cx*** |
| ***Aphanizomenon skujae*** | ***Aphanizomenon gracile cx*** |
| ***Aphanizomenon yezoense*** | ***Aphanizomenon flosaquae cx*** |
| *Aphanocapsa delicatissima* | *Aphanocapsa delicatissima* |
| *Aphanocapsa holsatica* | *Aphanocapsa holsatica* |
| *Aphanocapsa planctonica* | *Aphanocapsa planctonica* |
| *Asterionella formosa* | *Asterionella formosa* |
| *Aulacoseira alpigena* | *Aulacoseira alpigena* |
| *Aulacoseira ambigua* | *Aulacoseira ambigua* |
| *Aulacoseira distans* | *Aulacoseira distans* |
| ***Aulacoseira granulata*** | ***Aulacoseira granulata cx*** |
| ***Aulacoseira islandica*** | ***Aulacoseira islandica*** |
| ***Aulacoseira italica*** | ***Aulacoseira italica cx*** |

| | |
|---|---|
| ***Aulacoseira muzzanensis*** | ***Aulacoseira granulata cx*** |
| ***Aulacoseira subarctica*** | ***Aulacoseira italica cx*** |
| ***Botryococcus braunii*** | ***Botryococcus spp. cx*** |
| ***Botryococcus terribilis*** | ***Botryococcus spp. cx*** |
| *Ceratium furcoides* | *Ceratium furcoides* |
| *Ceratium hirundinella* | *Ceratium hirundinella* |
| *Ceratium rhomvoides* | *Ceratium rhomvoides* |
| *Chlamydocapsa planctonica* | *Chlamydocapsa planctonica* |
| *Chroococcus minutus* | *Chroococcus minutus* |
| *Chrysidiastrum catenatum* | *Chrysidiastrum catenatum* |
| *Chrysococcus cordiformis* | *Chrysococcus cordiformis* |
| *Chrysococcus ornatus* | *Chrysococcus ornatus* |
| *Chrysosphaerella longispina* | *Chrysosphaerella longispina* |
| *Closterium acutum* | *Closterium acutum* |
| *Coelastrum astroideum* | *Coelastrum astroideum* |
| *Coelastrum cambricum* | *Coelastrum cambricum* |
| *Coelastrum microporum* | *Coelastrum microporum* |
| *Coelastrum sphaericum* | *Coelastrum sphaericum* |
| *Coelomoron pusillum* | *Coelomoron pusillum* |
| *Crucigenia tetrapedia* | *Crucigenia tetrapedia* |
| ***Cryptomonas curvata*** | ***Cryptomonas spp. cx*** |
| ***Cryptomonas erosa*** | ***Cryptomonas spp. cx*** |
| ***Cryptomonas marssonii*** | ***Cryptomonas spp. cx*** |
| *Cuspidothrix issatschenkoi* | *Cuspidothrix issatschenkoi* |
| *Cyanodictyon imperfectum* | *Cyanodictyon imperfectum* |
| *Cyanodictyon planctonicum* | *Cyanodictyon planctonicum* |
| *Cyanodictyon reticulatum* | *Cyanodictyon reticulatum* |
| *Cyclotella meneghiniana* | *Cyclotella meneghiniana* |
| *Cyclotella radiosa* | *Cyclotella radiosa* |
| *Cyclotella stelligera* | *Cyclotella stelligera* |
| *Desmodesmus armatus* | *Desmodesmus armatus* |
| *Desmodesmus opoliensis* | *Desmodesmus opoliensis* |
| *Desmodesmus subspicatus* | *Desmodesmus subspicatus* |
| *Diatoma tenuis* | *Diatoma tenuis* |
| *Dimorphococcus lunatus* | *Dimorphococcus lunatus* |
| *Dinobryon bavaricum* | *Dinobryon bavaricum* |
| *Dinobryon crenulatum* | *Dinobryon crenulatum* |
| *Dinobryon divergens* | *Dinobryon divergens* |
| *Dinobryon sertularia* | *Dinobryon sertularia* |
| *Dinobryon sociale* | *Dinobryon sociale* |
| ***Dolichospermum affine*** | ***Dolichospermum spp. cx*** |
| ***Dolichospermum crassum*** | ***Dolichospermum spp. cx*** |
| ***Dolichospermum curvum*** | ***Dolichospermum spp. cx*** |
| ***Dolichospermum flosaquae*** | ***Dolichospermum spp. cx*** |
| ***Dolichospermum fuscum*** | ***Dolichospermum spp. cx*** |
| ***Dolichospermum lemmermannii*** | ***Dolichospermum spp. cx*** |
| ***Dolichospermum macrosporum*** | ***Dolichospermum spp. cx*** |
| ***Dolichospermum mendotae*** | ***Dolichospermum spp. cx*** |
| ***Dolichospermum mucosum*** | ***Dolichospermum spp. cx*** |
| ***Dolichospermum sigmoideum*** | ***Dolichospermum spp. cx*** |
| ***Dolichospermum smithii*** | ***Dolichospermum spp. cx*** |
| ***Dolichospermum solitarium*** | ***Dolichospermum spp. cx*** |
| ***Dolichospermum viguieri*** | ***Dolichospermum spp. cx*** |
| *Euglenaformis proxima* | *Euglena proxima* |
| *Eunotia zasuminensis* | *Eunotia zasuminensis* |
| *Fragilaria crotonensis* | *Fragilaria crotonensis* |
| *Gloeotrichia echinulata* | *Gloeotrichia echinulata* |
| *Golenkinia radiata* | *Golenkinia radiata* |

| | |
|---|---|
| *Gonyostomum latum* | *Gonyostomum latum* |
| *Gonyostomum semen* | *Gonyostomum semen* |
| ***Gymnodinium fuscum*** | ***Gymnodinium spp. cx*** |
| ***Gymnodinium uberrimum*** | ***Gymnodinium spp. cx*** |
| *Gyrodinium helveticum* | *Gyrodinium helveticum* |
| *Gyromitus cordiformis* | *Gyromitus cordiformis* |
| ***Hariotina reticulata*** | ***Pediastrum reticulatum cx*** |
| ***Lacunastrum gracillimum*** | ***Pediastrum duplex cx*** |
| ***Lemmermannia komarekii*** | ***Tetrastrum komarekii*** |
| *Limnococcus limneticus* | *Limnococcus limneticus* |
| *Mallomonas akrokomos* | *Mallomonas akrokomos* |
| *Mallomonas caudata* | *Mallomonas caudata* |
| *Mallomonas punctifera* | *Mallomonas punctifera* |
| *Mallomonas tonsurata* | *Mallomonas tonsurata* |
| *Melosira varians* | *Melosira varians* |
| *Merismopedia warmingiana* | *Merismopedia warmingiana* |
| *Micractinium pusillum* | *Micractinium pusillum* |
| ***Microcystis aeruginosa*** | ***Microcystis aeruginosa cx*** |
| ***Microcystis botrys*** | ***Microcystis aeruginosa cx*** |
| *Microcystis flos-aquae* | *Microcystis flos-aquae* |
| ***Microcystis novacekii*** | ***Microcystis aeruginosa cx*** |
| *Microcystis viridis* | *Microcystis viridis* |
| *Microcystis wesenbergii* | *Microcystis wesenbergii* |
| *Monoraphidium dybowskii* | *Monoraphidium dybowskii* |
| *Mucidosphaerium pulchellum* | *Mucidosphaerium pulchellum* |
| *Nitzschia holsatica* | *Nitzschia holsatica* |
| *Oocystis borgei* | *Oocystis borgei* |
| *Oscillatoria tenuis* | *Oscillatoria tenuis* |
| *Pandorina morum* | *Pandorina morum* |
| *Parvodinium goslaviense* | *Parvodinium goslaviense* |
| *Parvodinium umbonatum* | *Parvodinium umbonatum* |
| ***Pediastrum boryanum*** | ***Pseudopediastrum boryanum*** |
| ***Pediastrum duplex*** | ***Pediastrum duplex cx*** |
| *Peridinium cinctum* | *Peridinium cinctum* |
| *Peridinium willei* | *Peridinium willei* |
| ***Plagioselmis nannoplanctica*** | ***Rhodomonas lacustris cx*** |
| *Planktolyngbya limnetica* | *Planktolyngbya limnetica* |
| *Planktothrix agardhii* | *Planktothrix agardhii* |
| *Pseudanabaena limnetica* | *Pseudanabaena limnetica* |
| *Pseudanabaena mucicola* | *Pseudanabaena mucicola* |
| *Pseudogoniochloris tripus* | *Pseudogoniochloris tripus* |
| ***Pseudopediastrum boryanum*** | ***Pseudopediastrum boryanum*** |
| *Pseudosphaerocystis lacustris* | *Pseudosphaerocystis lacustris* |
| *Radiocystis geminata* | *Radiocystis geminata* |
| ***Rhizosolenia longiseta*** | ***Urosolenia longiseta*** |
| ***Rhodomonas lacustris*** | ***Rhodomonas lacustris cx*** |
| *Scenedesmus ellipticus* | *Scenedesmus ellipticus* |
| *Scenedesmus obtusus* | *Scenedesmus obtusus* |
| *Scenedesmus quadricauda* | *Scenedesmus quadricauda* |
| *Skeletonema potamos* | *Skeletonema potamos* |
| *Snowella septentrionalis* | *Snowella septentrionalis* |
| *Sphaerocystis schroeteri* | *Sphaerocystis schroeteri* |
| *Spondylosium planum* | *Spondylosium planum* |
| *Staurastrum anatinum* | *Staurastrum anatinum* |
| *Staurastrum luetkemuelleri* | *Staurastrum luetkemuelleri* |
| *Staurastrum paradoxum* | *Staurastrum paradoxum* |
| *Stauridium privum* | *Stauridium privum* |
| *Stauridium tetras* | *Stauridium tetras* |

| | |
|---|---|
| *Staurodesmus dejectus* | *Staurodesmus dejectus* |
| *Staurodesmus mucronatus* | *Staurodesmus mucronatus* |
| *Stephanodiscus binderanus* | *Stephanodiscus binderanus* |
| *Stephanodiscus hantzschii* | *Stephanodiscus hantzschii* |
| *Stephanodiscus rotula* | *Stephanodiscus rotula* |
| *Stichogloea doederleinii* | *Stichogloea doederleinii* |
| ***Synedra ulna*** | ***Ulnaria ulna cx*** |
| ***Synura petersenii*** | ***Synura spp. cx*** |
| ***Synura uvella*** | ***Synura spp. cx*** |
| *Tabellaria fenestrata* | *Tabellaria fenestrata* |
| *Tabellaria flocculosa* | *Tabellaria flocculosa* |
| *Tetraedron caudatum* | *Tetraedron caudatum* |
| *Tetraedron minimum* | *Tetraedron minimum* |
| *Trachelomonas crebea* | *Trachelomonas crebea* |
| *Trachelomonas hispida* | *Trachelomonas hispida* |
| *Trachelomonas intermedia* | *Trachelomonas intermedia* |
| *Trachelomonas planctonica* | *Trachelomonas planctonica* |
| *Trachelomonas volvocina* | *Trachelomonas volvocina* |
| *Trachelomonas volvocinopsis* | *Trachelomonas volvocinopsis* |
| ***Ulnaria delicatissima*** | ***Ulnaria ulna cx*** |
| ***Ulnaria ulna*** | ***Ulnaria ulna cx*** |
| *Urosolenia eriensis* | *Urosolenia eriensis* |
| *Westella botryoides* | *Westella botryoides* |
| *Woronichinia naegeliana* | *Woronichinia naegeliana* |

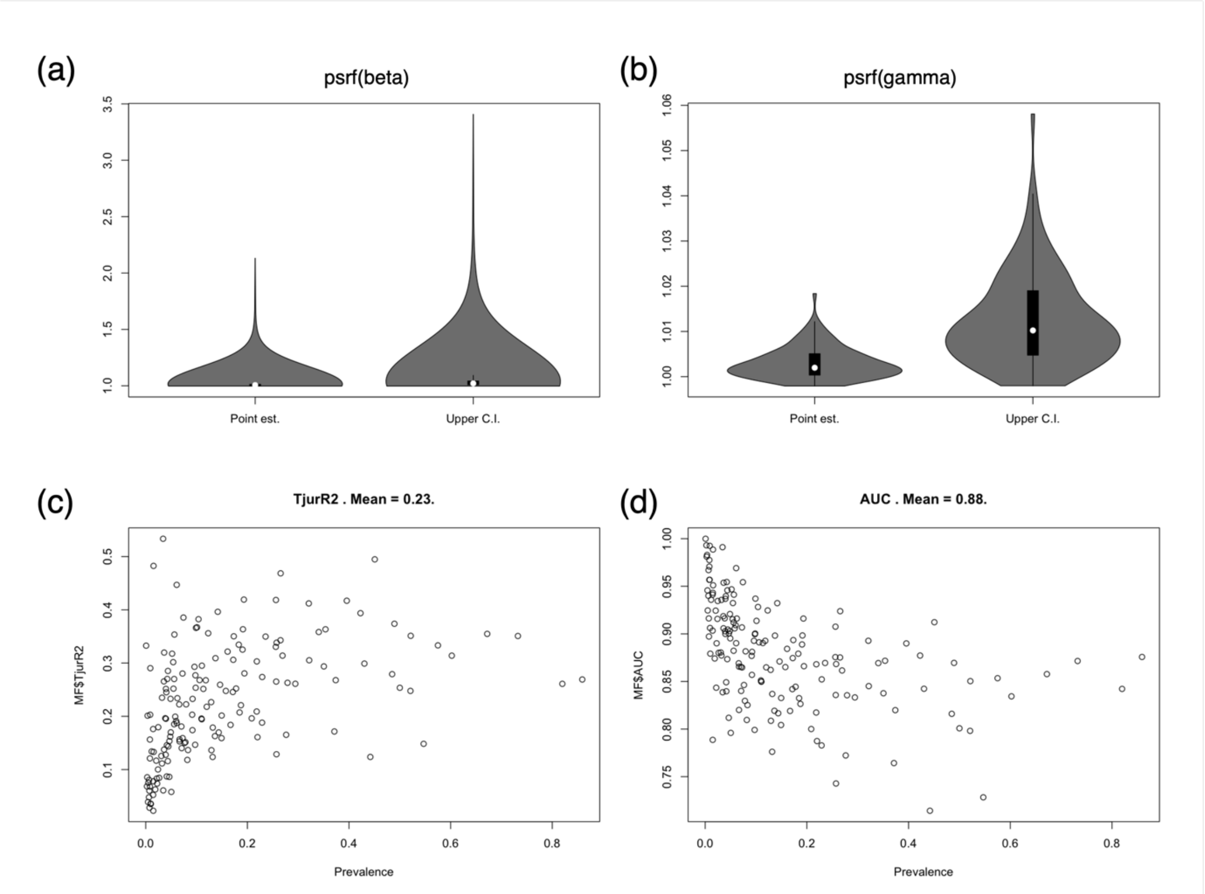


**Figure S1:** Diagnostics for MCMC convergence via potential scale reduction factor for (a) beta parameters (species-environment), (b) gamma parameters (trait-environment) and model fit with (c) species specific Tjur $R^2$ and (d) AUC values over species prevalence.

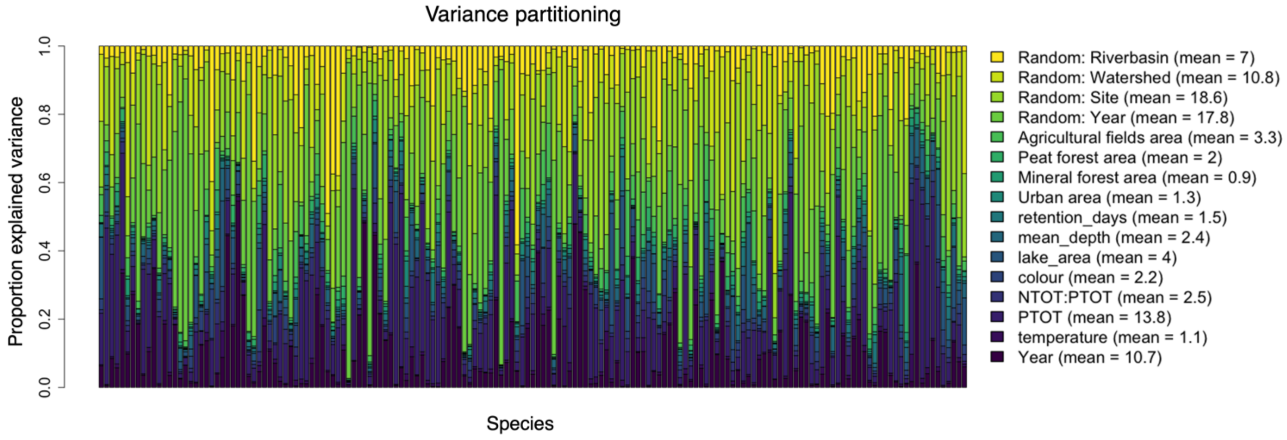


**Figure S2:** Results on variance partitioning. Variation in species occurrences is partitioned into responses to fixed and random effects. The bar-plot shows species-specific results whereas the legend shows averages over all species.

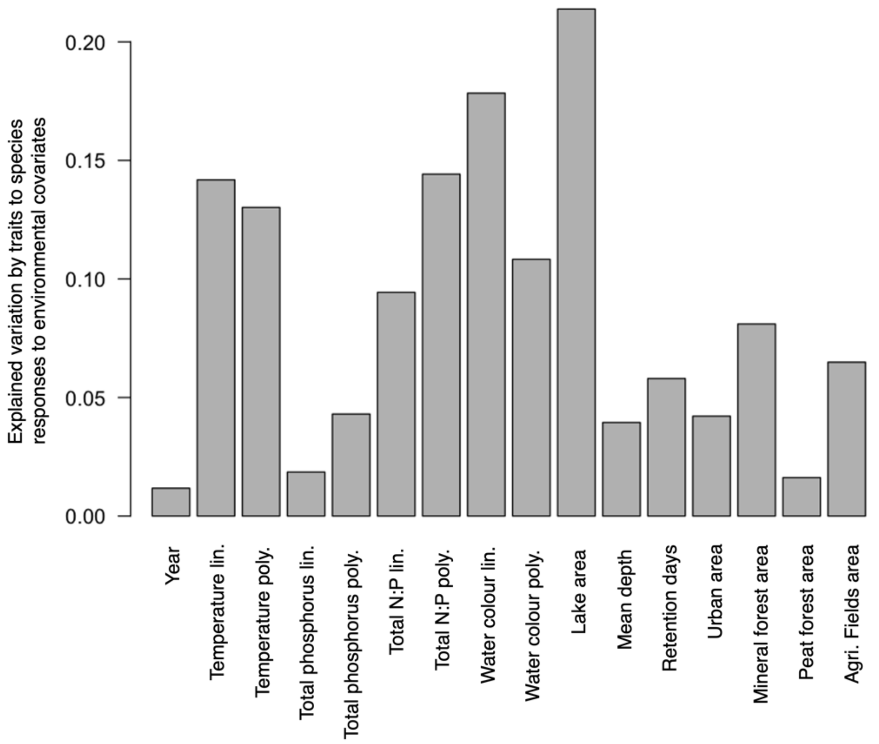


**Figure S3:** Proportion of explained variation by traits among the species responses to environmental covariates

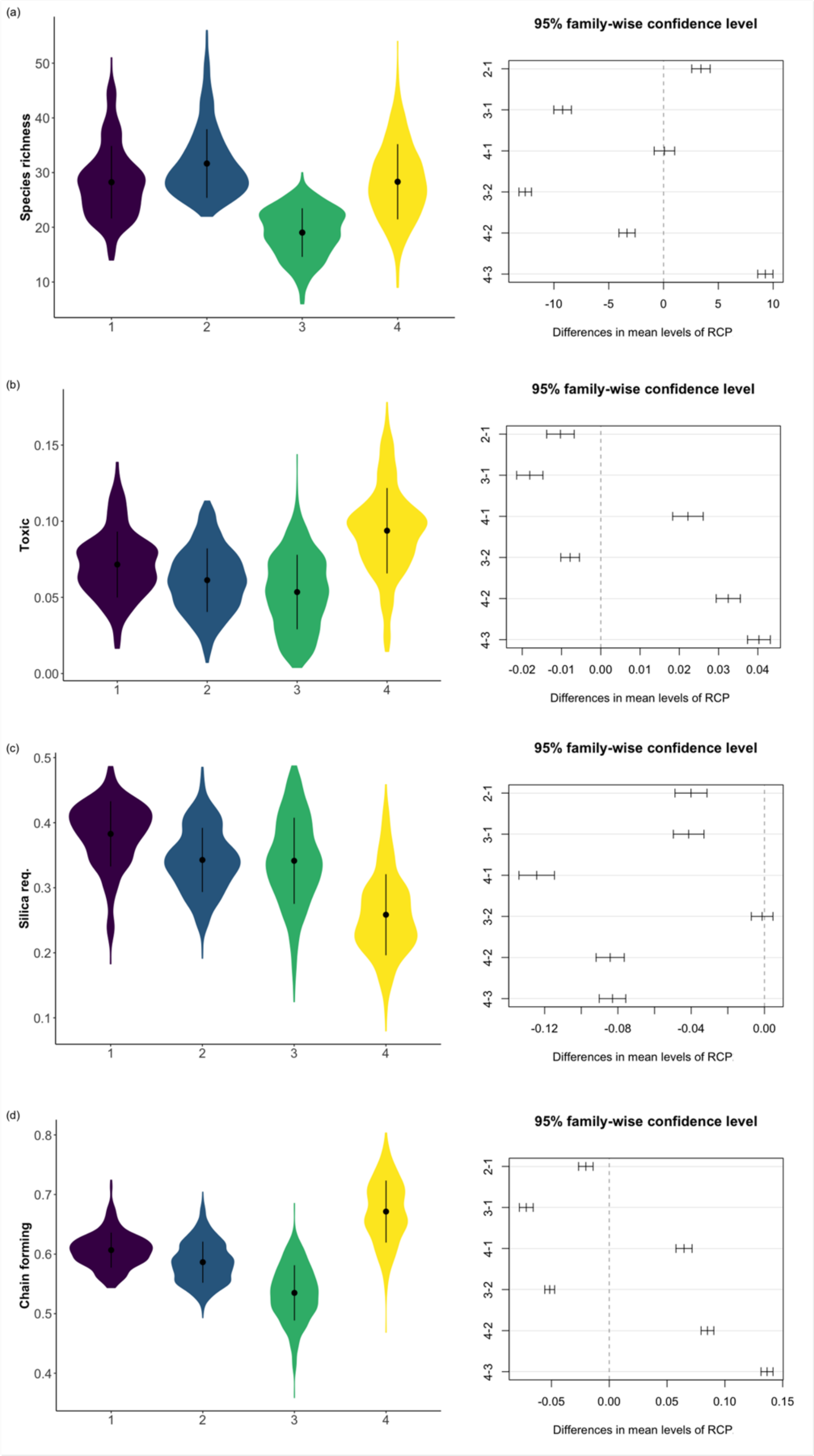

(a)
Species richness
95% family-wise confidence level
Differences in mean levels of RCP
(b)
Toxic
95% family-wise confidence level
Differences in mean levels of RCP
(c)
Silica req.
95% family-wise confidence level
Differences in mean levels of RCP
(d)
Chain forming
95% family-wise confidence level
Differences in mean levels of RCP

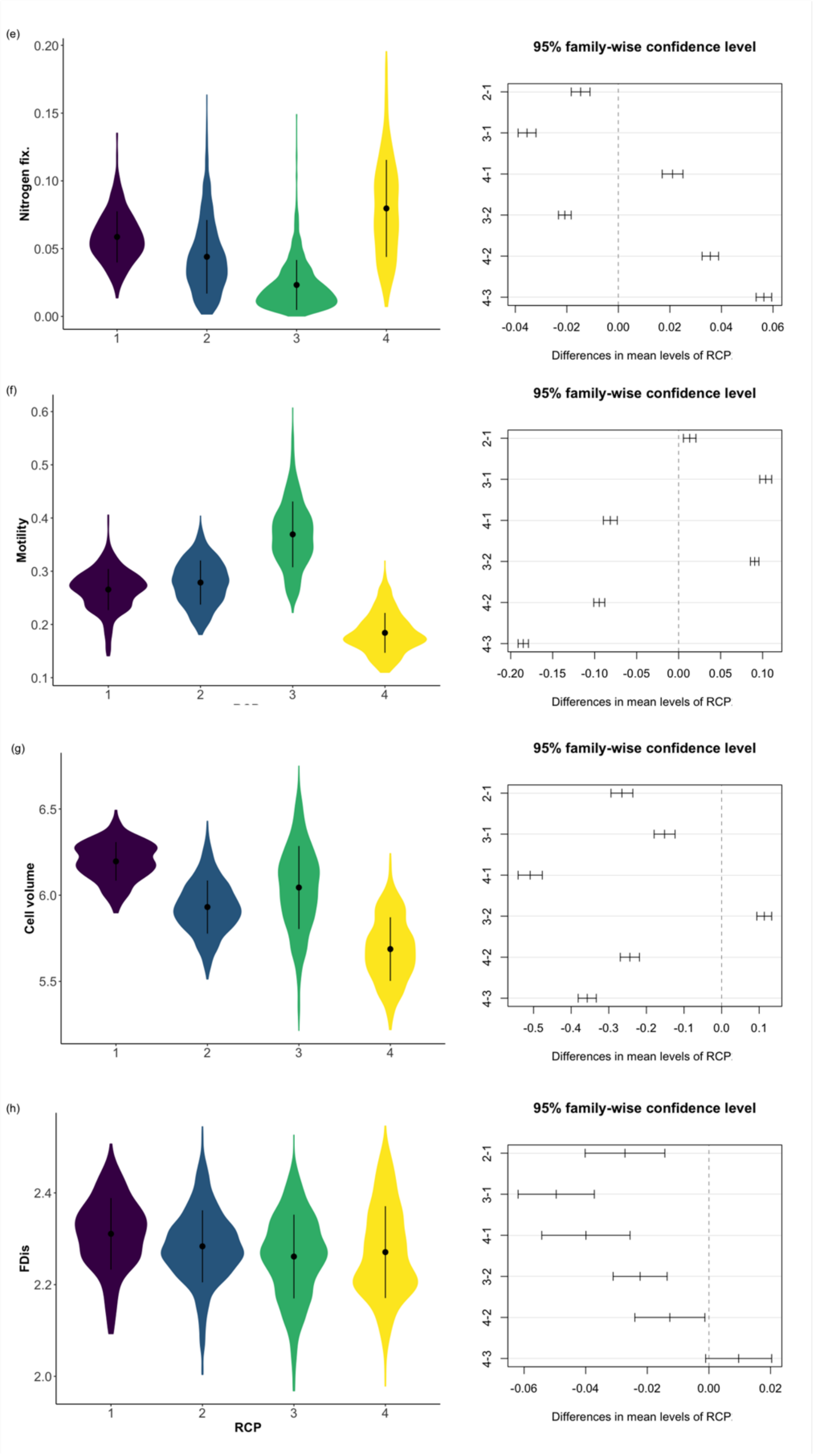


**Figure S4:** Illustration of community characteristics at the four Regions of Common Profile (RCP) as shown in main manuscript, here highlighting statistical differences between means of RCPs using Tukey's post hoc test for (a) species richness (b-g) community weighted trait values, (h) functional diversity displayed as Functional Dispersion FDis. Colours of RCPs are the same as in Fig. 3.

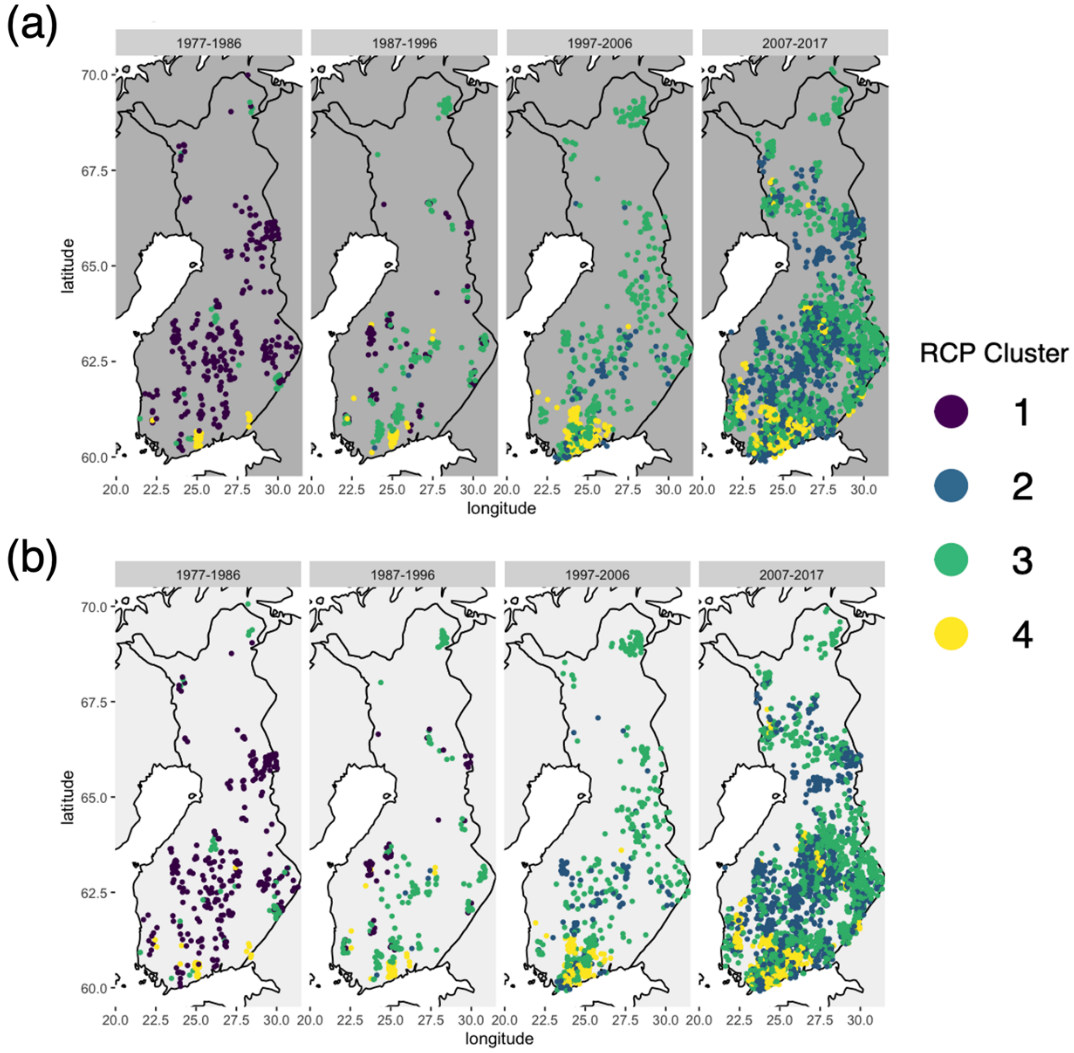


**Figure S5:** Regions of common profile based on optimal clustering of predicted species compositions for (a) model including the full species list and (b) model including the adjusted species complex list from Table S1. Data are displayed in aggregations of 10 years to highlight the temporal aspect of change. To avoid overlapping data points, the data are plotted with jitter of 0.2 degree for latitude and longitude.
